# Rotating Letters in the Mind’s Eye: Behavioral and electro-cortical associations with 3D Mental-Rotation Ability

**DOI:** 10.64898/2026.05.11.724360

**Authors:** Rozemun Khan, Stamatia Bekiari, Beerend Hierck, Daniela Salvatori, J. Leon Kenemans

## Abstract

Mental rotation in 3D is a key cognitive skill involving dynamic spatial transformations, for which pronounced individual differences have been documented. Here we ask whether individual differences in 3D abilities can be explained by analogous differences in 2D abilities. 3D mental-rotation was assessed by the Vandenberg & Kruse Mental Rotation Test (3D-MRT) and examined for association with performance and underlying electrocortical mechanisms during a 2D letter rotation task.

Participants (N=40) first completed the MRT and then performed a computerized 2-D letter rotation task in which they had to identify whether letters were oriented in a standard or a mirrored direction (parity judgment) when rotated at 0°, 60°, 120°, and 180° while EEG was recorded.

Reaction times (RTs) and error rates increased with angular disparity. The angular disparity effect on RT was smaller for mirrored letters. Low, relative to high, 3D-MRT scoring participants showed more pronounced accuracy declines at higher rotation angles.

An EEG Event Related Potential (ERP) known as the Rotation-Related Negativity (RRN) became more pronounced with increasing angular disparity. High 3D-MRT scores were associated with a stronger RRN response at central-parietal sites. In addition, the ERP-P3b wave was more pronounced at central-parietal sites for low 3D-MRT scorers, independent of angular disparity. It is concluded that 3D rotational ability is positively associated with 2D mental rotation performance, and more strongly with enhanced recruitment of neural visual-spatial cortical representations than with enhanced recruitment of more general cognitive resources.

## Introduction

Visuospatial abilities allow for understanding, reasoning about, and remembering spatial relationships among spaces and objects (Lohman, 1996; Hegarty & Waller, 2005). They represent an overarching cognitive skill composed of several subcomponents, including the ability to visualize and mentally manipulate objects in space, navigate through environments, and interpret relationships between different spatial configurations (Hegarty et al., 2006; Montello, 1993). These skills are extremely varied across different individuals and are crucially important in domains that require spatial reasoning, such as engineering, architecture, aviation, medicine (including veterinary), and anatomy education. This is particularly relevant as such fields increasingly rely on three-dimensional technologies like eXtended Reality (XR), which require learners to interpret complex three-dimensional structures and their spatial interrelations (Wai, Lubinski, & Benbow, 2009).

A subcomponent of visuospatial ability is mental rotation which involves the transformation of an internal representation of an object to assess features regarding its identity or orientation (Shepard & Metzler, 1971; Just & Carpenter, 1985). Mental rotation is a skill linked to navigation, technical problem-solving, and a range of STEM professions. Some researchers have indicated that individuals with stronger spatial reasoning are more likely to pursue and excel in STEM disciplines, and that training of visuospatial skills has been shown to enhance STEM learning outcomes (Wai, Lubinski, & Benbow, 2009).

A way to categorize individual proficiency in 3D mental rotation (MR) is through the Mental Rotation Test (MRT) which was developed by Vandenberg and Kuse (1978) as a standardized, paper-and-pencil adaptation of the original Shepard and Metzler (1971) mental rotation paradigm. Participants are asked to compare pairs of three-dimensional figures and decide if they are the same object rotated in space or different objects. This task requires the ability to visualize the rotation of objects in the mind’s eye and match them accordingly (Shepard & Metzler, 1971; Voyer et al., 1995).

Research conducted by Bogomolova et al. (2020) and others has expanded on the relationship between individual 3D-MR ability and performance on tasks involving spatial visualization. They found that individuals with lower, compared to those with higher, 3D-MR ability benefitted more when learning anatomical structures using stereoscopic (3D) rather than monoscopic (2D) renderings of these 3D structures (Bogomolova et al., 2020). These findings suggests that dimensionality interacts with 3D-MR ability to impact how much individuals can benefit from and process complex spatial information and educational content (Chikha et al., 2021). However, it remains unclear how individual differences in 3D-MR ability relates to processing of 2D rotated figures.

Understanding rotation in 2D space, here described as 2D rotation, is important because it is essential to characterize how individuals perform *rotation along a planar axis* without additional depth cues, perspective changes, and object-complexity demands inherent to more complex 3D stimuli. In 2D tasks, such as classic letter rotation paradigms, participants mentally rotate planar objects to judge whether they are in standard or mirrored configuration (Cooper, 1975). Typically, performance in terms of speed and accuracy is reduced with increasing rotation demand, i.e., increasing angular disparity of the letter from the canonical orientation. This raises an open question: Do high 3D-MR scorers also excel at 2D rotation? If so, we would expect steeper declines in performance with increasing angular disparity for low, relative to high 3D-MR performers.

Previous studies have identified a characteristic EEG event-related potential (ERP) known as the rotation-related negativity (RRN). The RRN typically emerges at central-parietal scalp sites around 400 ms after stimulus onset and can extend to approximately 700–800 ms (Wijers et al., 1989; Farah & Peronnet, 1989; Riečanský & Jagla, 2008). The amplitude of the RRN varies as a function of angular disparity; it becomes more negative with greater angular disparities (Heil, 2002; ter Horst et al, 2010; Zhao et al., 2018). The work of Heil (2002; Heil & Rolke, 2002) also demonstrates that the RRN is specific for mental rotation, and is not elicited by, e.g., additional pixel noise or increased difficulty of (letter) discrimination. This makes the RRN a valuable marker to explore how individuals differ in the neural substrate of 2D-MR. The specific association between RRN and the mental-rotation process is consistent with the notion that RRN reflects the increased recruitment of visual-spatial cortical representations (Gardony et al., 2017); this will be referred to as the representation perspective.

However, the RRN evolves in globally the same time window as does the P3b and also has a similar central-parietal distribution (see Heil; Heil & Rolke, 2002). The P3b is a positive deflection thought to reflect updating of (working) memory representations in response to task-relevant stimuli, with larger amplitudes indexing greater allocation of attentional and cognitive resources (Polich, 2007). Critically, as demonstrated by Bajric et al. (1999), in contrast to RRN in a mental-rotation-parity-judgement task, the P3b increases in positivity with increasing angular disparity. This fits the notion that P3b scales with allocation of cognitive resources, which we refer to as the resource perspective. As Bjaric et al. note, theoretically, the increase with increasing angular disparity in P3b positivity and in RRN negativity may cancel each other out. However, the RRN literature on neurotypical individuals shows that the signal becomes increasingly negative with rotation angle, indicating that the RRNs negative effects dominate the P3b’s positive effects.

When considering neurotypical individuals with differing levels of 3D-MR ability, a number of alternative hypotheses can be formulated as to the differential effects of angular disparity. First, individual differences may be absent if individuals recruit visuospatial representations (reflected by RRN) and cognitive resources (reflected in P3b) to the same degree, or if both components increase proportionally within a given group (e.g., with enhanced recruiting in high 3D-MR scorers of both representations and global resources), cancelling out net differences. Second, high 3D-MRT scorers may show more negative angular-disparity effects (larger RRN modulation) if representation recruitment dominates over resource allocation relative to low 3D-MRT individuals. This pattern would especially emerge when in high 3D-MRT individuals, representations are more strongly recruited, and/or for low 3D-MR individuals resources more strongly (e.g., because the task is experienced as more difficult for low 3D scorers). Third, this pattern would be reversed if high 3D-MRT scorers rely more on global cognitive resources and less on visuospatial representations, resulting in a stronger P3b and less negative RRN for high-3D scorers compared to low-3D scorers.

The present study therefore investigates whether individual differences in 3D-MR ability are associated with distinct behavioral and RRN modulation patterns during a 2D letter rotation task. Specifically, we predict that 2D-rotation-angle-dependent declines in performance are associated with low 3D-MRT scores. As to the electrocortical level, we predict that high 3D-MRT scorers will show larger RRN modulation with angular disparity compared to low 3D-MRT scorers, reflecting greater reliance on visuospatial representations relative to global cognitive resources (second hypothesis above).

## Methods

### 2.1. Participants

Sample size was determined in reference to previous ERP work using comparable mental rotation paradigms (Riečanský & Jagla, 2008; N=14). Because the present study included an additional between-subject factor related to individual differences in MR ability, recruitment aimed to approximately double this sample size. Fifty-two participants were recruited. Participants received either 20 euros for 2 hours or course credit. Twelve participants had to be excluded because of: early termination during data collection, data loss, insufficient data quality and/or insufficient performance on the behavioral task. Ultimately, 40 participants (24 F, 16 M), between 19 and 50 years of age (mean 26.22, ± 6.25 years) were considered for data analyses. Informed consent was obtained from all participants, and the study procedures were approved by the Ethics Board of the Utrecht University Faculty of Social and Behavioral Sciences (file number 24-0457).

### 2.2. 3D Mental Rotation Test

Participants were administered a 24-item Mental Rotation Test (3D-MRT; Vandenberg & Kruse, 1978). The questions were structured with a template 3D figure on the left and four 3D figures on the right (Figure 1). The participant needed to decide which two of the four objects on the right were identical (except for being rotated) to the template on the left. The test results served as an estimate (range 0-24) for individual MR ability.

**Figure 1.**
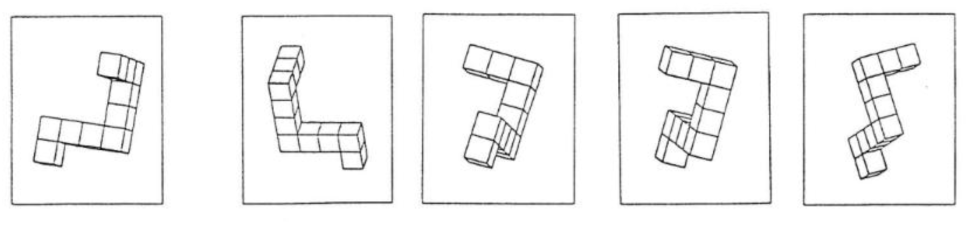
Example item from the Vandenberg and Kruse Mental Rotation Test (1978). The figure on the left serves as the reference stimulus. Participants are required to identify the two figures, among the four options, that are identical to the reference except for being rotated. One point is awarded for each item only when both correct choices are selected. Partially correct answers were not awarded. In the example, the most left and the third option on the right match the reference.

### 2.3. Experimental procedure

Participants were seated in an electrically shielded room for the task. They read instructions about the main task and the 3D-MR test and signed a consent form when done. Before attempting the 3D-MR test, they were asked to do four practice questions. The MRT was then administered as a pen and paper version of the test, which the participants had 10 minutes to complete. The participants were also reminded when they had 5 minutes left and then once again when they had 2 minutes left. At the end of the 10 minutes, the test was collected from the participant.

Following the 3D-MR test the participant was set up for EEG recording with their head positioned on a chin rest such that there was a distance of 60 cm from the nasion to the computer monitor. Familiarization with the letters and their rotated forms, and a 30-trial practice letter rotation task followed. Once practice performance reached a minimum of 80% accuracy the participant could proceed with the main task. During the practice task, participants received automatic feedback on their selection (‘S’ for standard orientation or ‘M’ for mirrored orientation) for each trial. The screen displayed “correct” or “incorrect”, and by the end of the practice task the participant received a score. This feedback on accuracy was not present during the experimental phase of the task.

### 2.4. Task & Stimuli

A total of 7 letters were used in the main letter rotation task: F, G, J, R, Q, P, L. The letters were presented in white on a black background. The letters were created in Adobe Photoshop at a height of 1.04 cm, which corresponds to a visual angle of ≈ 0.99° at a viewing distance of 60 cm programmed in PsychoPy2 (version 2024.2.4; Peirce et al., 2019). Stimuli were presented centrally and displayed in either standard or mirrored form (flipped along the y-axis) in Arial font.

The stimuli were scaled proportionally to fit the display. These letters were presented at 4 different orientations: 0°, 60°, 120°, 180°. At the angles of 60° and 120° the letters were presented in both clockwise (C) and counterclockwise (CC) direction (Figure 2). Letters were presented in standard (canonical) and mirrored (reversed) conditions.

**Figure 2.**
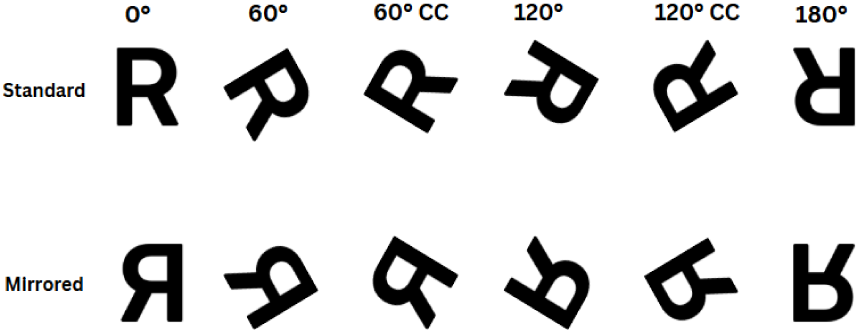
Examples of the orientations of standard and mirrored letters used as stimuli in the present study.

Participants needed to press ‘S’ on their keyboard when they saw a letter in standard condition and ‘M’ when the letter was mirrored. Trials started with a fixation cross for a variable interval of 900-1100 ms, followed by stimulus presentation (250 ms), and ended within a time window for responses which was fixed at 1900 ms, irrespective of response latency (Figure 3). Participants were instructed to keep their eyes on the fixation cross throughout the task, and their fingers on the two keys (S and M) on the keyboard in front of them, which were taped with S and M written with black marker on top to distinguish from other keys. They had to respond as quickly and accurately as possible to determine the parity of the letter on the screen.

**Figure 3:**
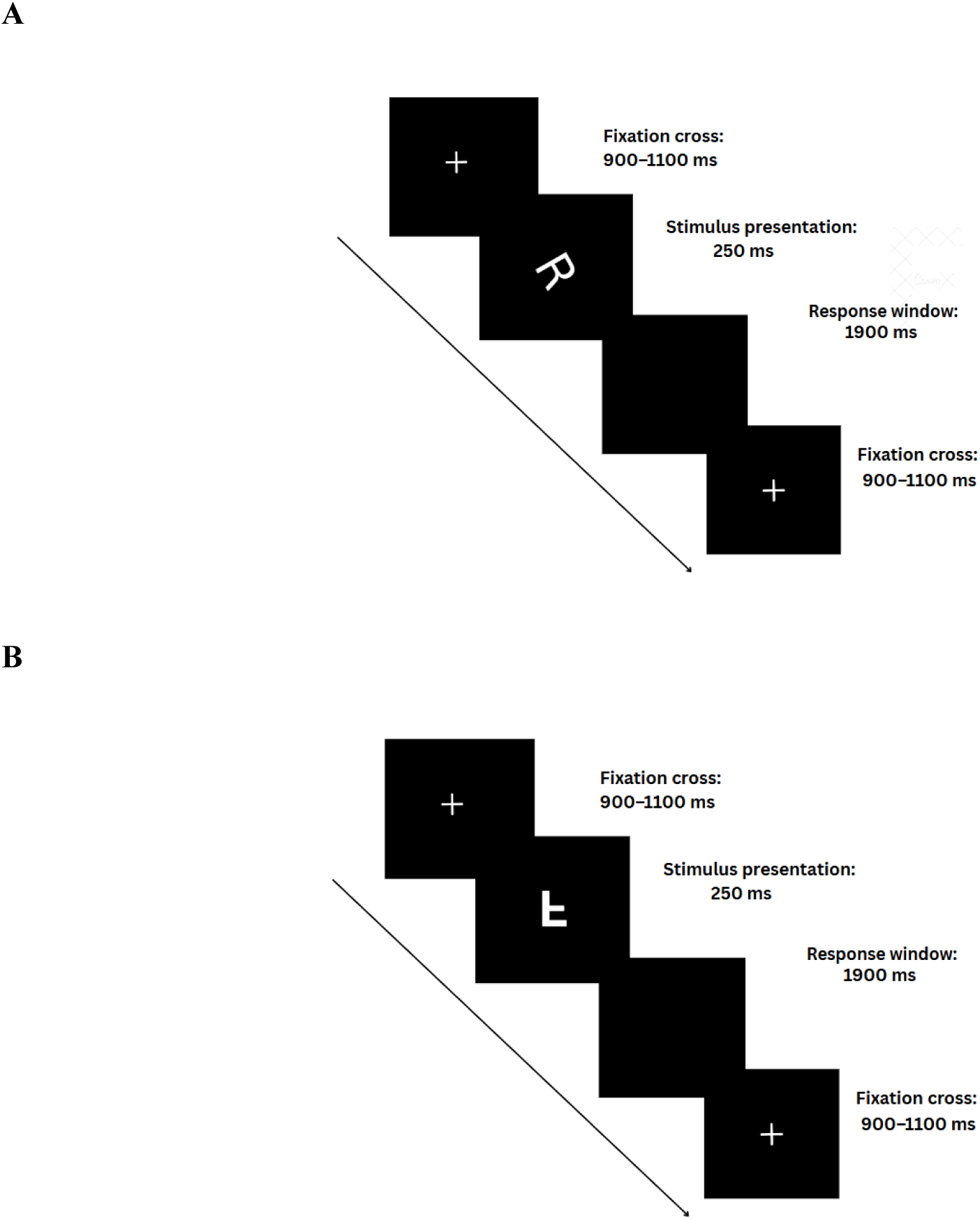
Example Trials. **A)** shows an example of a trial where the stimulus appears for 250 ms in an Standard orientation rotated 60 degrees. During the 250 ms stimulus presentation time window, no responses are registered, after 250 ms the stimulus disappears and the participant would need to select ‘S’ in the 1900 ms time window for the response to be registered as correct. **B)** shows a different trial with the stimulus now appearing in 180 degrees Mirrored orientation. The participant would need to press ‘M’ in the 1900 ms response window for the response to be registered as correct.

The experiment had 560 trials split between 5 blocks with each block having a total of 112 trials (7 letters × 2 versions (parity) × 4 orientations, including CC conditions for 60 and 120), including repeats of combinations. Between blocks the participants could take a short break and manually continue with the experiment whenever they were ready by pressing any key on the keyboard.

The order of the trial presentation of letters was randomized separately for each participant beforehand. Trial order was pseudorandomized within predefined constraints to ensure balanced presentation of all conditions and to reduce order effects. That is, no letter was presented more than twice consecutively, and no stimulus version (standard or mirrored) appeared more than four times in a row.

### 2.5. Electrophysiological recordings

EEG was recorded using a 64-channel BioSemi ActiveTwo system (BioSemi B.V., The Netherlands). Electrode position layout followed the 10/10 system. Six external electrodes were used. Four were used for ocular correction: two were placed above and below the left eye to capture vertical saccades and eyeblinks, and two placed at the outer canthi of the left and right eye to capture horizontal saccades. Two additional electrodes were also placed on the right and left mastoid bones and were used for offline re-referencing. Data were collected at a sampling rate of 2048 Hz, using an 418 Hz low-pass filter, and with electrode impedances below 50 kΩ.

### 2.6. Performance data processing

Three types of responses were recorded: correct, incorrect, or missed response. Misses occurred from 0.89% to 1.94 % across conditions and participants. Trials with reaction times 250 ms or shorter were not recorded. Incorrect responses were used to calculate accuracy percentages per condition. Mean reaction times per condition were based on correct responses.

### 2.7. EEG data pre-preprocessing

Pre-processing of the EEG data was done offline in Brain Vision Analyzer (BVA) Software Version 2.1 (Brain Products, Munich, Germany). Data were band-pass filtered between 0.01 and 30 Hz, resampled to 256 Hz, and re-referenced to the averaged mastoid electrodes.

Next, the data were segmented into trial epochs of −600 ms before stimulus onset up to 2100 ms. A first round of artifact rejection was applied to the channels AFz, CPz, Cz, FCz, Fpz, Fz, Oz, P3, P4, POz, and Pz, including: voltage step(s) exceeding 50 μV between sample points, within-segment differences exceeding 200 µV, and absolute amplitudes exceeding ± 200 μV; also a low-activity criterion of 0.5 μV within 100 ms intervals was included to detect flat or near-flat segments. This was followed by Gratton-Coles ocular-artifact correction, in turn followed by another round of artifact rejection with the same parameters except for within-segment differences exceeding 100 µV, to eliminate residual ocular artifacts.

When calculating average ERPs, a criterion was set of there being at least 20 segments per each of the 8 condition combinations, in order for the data of that participant to be included. Participant-level ERPs were averaged separately for each of the eight conditions (0M, 0S, 60M, 60S, 120M, 120S, 180M, 180S; M = mirrored, S = standard). Baseline correction was applied using the 50 ms preceding stimulus onset.

In the average ERPs, rotation-related negativity (RRN) was quantified as the average amplitude in the 450–750-ms latency window. Although it was expected that RRN would be maximal at Pz (or CPz), values for more anterior and posterior analyses were also included to account for inter-individual differences as to the exact location of the maximum values; the complete array included six midline electrodes (FCz, Cz, CPz, Pz, POz, Oz). RRN values were extracted separately for each participant, parity, and orientation.

### 2.8. Statistical Analysis

Statistical analyses were conducted with R software, version 4.1.2 (R Core Team, 2021). For data preparation the tidyverse (Wickham et al., 2019) and readxl (Wickham & Bryan, 2024) packages were applied. Performance (RT, accuracy) and ERP values across letter angles were transformed to three polynomial-term trend scores (0-order, linear, quadratic), using R’s contr.poly() function. The polynomial contrasts were convoluted with the averages across and differences between Parity levels (standard, mirrored). This resulted in a 3×2 Orientation-trend scores x Parity contrast scores, which were in turn entered in R’s basic t test function to test against zero.

#### 2.8.1. Performance Measures

RT and accuracy were analyzed using the general approach outlined above. In a second step, associations between task performance and 3D-MRT scores were analyzed using the forward multiple regression procedure in SPSS (Version 30.0; IBM Corp., 2024). This involves the following steps: 1) compute all the correlations between the dependent variable 3D-MRT and the 12 independent variables (contrasts for Parity (2) x Orientation (3) x Measure (RT, accuracy)); 2) create a first regression model with the highest correlating independent variable as the predictor; 3) then add the second highest correlating independent variable; 4) evaluate whether both predictors are significant (at p < .05); 5) if they are not, then the first model is selected; if they are the third highest correlating variable can be added, and steps 4 and 5 are repeated. For confirmatory purposes the eventually selected model can also be evaluated (relative to the preceding, simpler model) in terms of adjusted R^2^ and Akaike and Bayes information criteria (AIC and BIC). This whole procedure can also be implemented by first evaluating all correlations (including interaction terms), and then conducting the subsequent regression analyses one by one, but the automated forward procedure is much more efficient, if only because a researcher does not need to first inspect multiple correlations.

#### 2.8.2. ERP: RRN/P3b

RRN/ P3b data were analyzed following the general approach, supplemented with the electrode factor. Electrode 0-order, linear, and quadratic trend scores (capturing anterior-posterior gradients) were computed using R’s `contr.poly()` function. This approach results in a test space consisting of 3 Orientation trends times 2 Parity scores times 3 Electrode trend scores, yielding 18 tests in total. To control for Type-I errors, the critical p value was 0.003 (0.05/18).

In a second step, all 9 combined Orientation and Electrode trend scores (averaged across Parity levels) were entered as predictors in a stepwise regression analysis with 3D-MRT score as dependent variable. This procedure aimed at probing both RRN and P3b spatial patterns across Orientation levels for the extent to which they significantly predicted 3D-MRT score. See the preceding section for a detailed explanation of this procedure.

### 2.9. ERP: Exploratory Analysis

To identify additional time points and scalp regions showing effects of mental rotation and individual differences in spatial ability, an exploratory analysis was conducted using a baseline-corrected cluster-based procedure (Chen et al., 2024). ERP amplitudes were computed across 24 electrodes organized into an 8 (anterior– posterior) × 3 (left–center–right) topographical grid: AF3-AFz-AF4; F3-Fz-F4; FC3-FCz-FC4; C3-Cz-C4; CP3-CPz-CP4; P3-Pz-P4; PO3-POz-PO4; O1-Oz-O2. Data were segmented into 100 ms non-overlapping time bins spanning 100–1100 ms post-stimulus, with a pre-stimulus baseline of −600 to 0 ms. Spatial structure was captured using orthogonal polynomial contrasts (0-order, linear, quadratic) along each axis, reducing spatial noise while preserving interpretable topographic gradients. Baseline (−600 to 0 ms; 7 bins) and (active) post-stimulus (100 to 1100 ms; 10 bins) periods were analyzed to establish empirical cluster size thresholds. The baseline alpha was set to .07 (0.05 times 10/7) to account for the slightly larger number of active relative to baseline bins and to identify the largest cluster expected by chance under the null hypothesis. Post-stimulus effects were tested using Bonferroni correction (α = .05/54 = .001) to account for multiple comparisons across 9 spatial contrasts × 3 orientation contrasts × 2 parity = 54 tests. With electrode main effects excluded, 45 tests remained, yielding a corrected p-value criterion of .05/45 = .001. Only clusters with consecutive significant bins (p < .001) exceeding the empirically derived baseline cluster threshold were considered statistically meaningful.

## 3. RESULTS

### 3.1. 3D-MRT Scores

Mental Rotation Test scores for the 40 participants ranged from 3 to 24 (*M* = 13.9, *SD* = 4.69, *Mdn* = 13, IQR = 5.25). A Shapiro-Wilk test confirmed that 3D-MRT scores were normally distributed, *W* = 0.98, *p* = .67 (Figure 4). For all statistical analyses, 3D-MRT score was treated as a continuous variable to preserve the full range of individual differences in spatial ability. However, for visualization purposes, in performance and waveform plots participants were grouped into Low (≤11; n=9), Medium (12-15; n=18), and High (≥16; n=13) spatial ability categories to illustrate patterns across different 3D-MRT levels. Alternative grouping schemes (e.g. tertile splits or median splits) would similarly produce unequal groups given the sample size and score distribution. The inclusion of a Medium group reflected clustering around the center of the distribution (e.g., several participants scored 12. These groupings were used solely for descriptive visualization and do not reflect the analytical approach.

**Figure 4.**
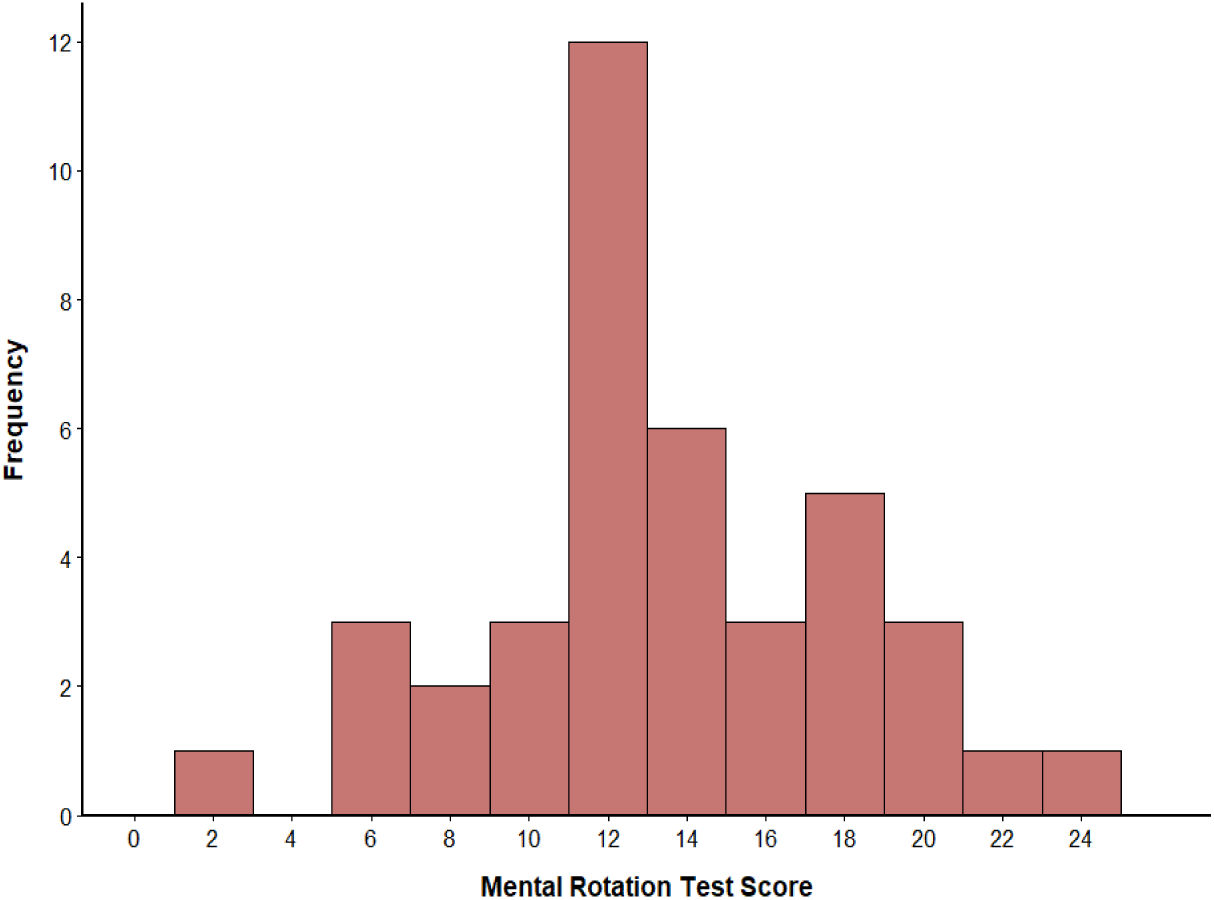
3D-MRT test score distribution. Bars represent 2-point bins spanning consecutive scores (e.g., the bar at 12 includes scores 12 and 13). Frequency indicates the number of participants per bin. Scores ranged from 3 to 24 (M = 13.9, SD = 4.69, Mdn = 13).

### 3.2. Performance

Reaction time increased with rotation angle in both standard and mirrored conditions (Figure 5). The statistical analysis revealed significant main effects of Parity (t(39) = 13.4, p < .001), Orientation (linear) and (quadratic), as well as interactions between Parity and the two Orientation trend effects (all ps < .001; smallest t(39) = 3.6).

**Figure 5:**
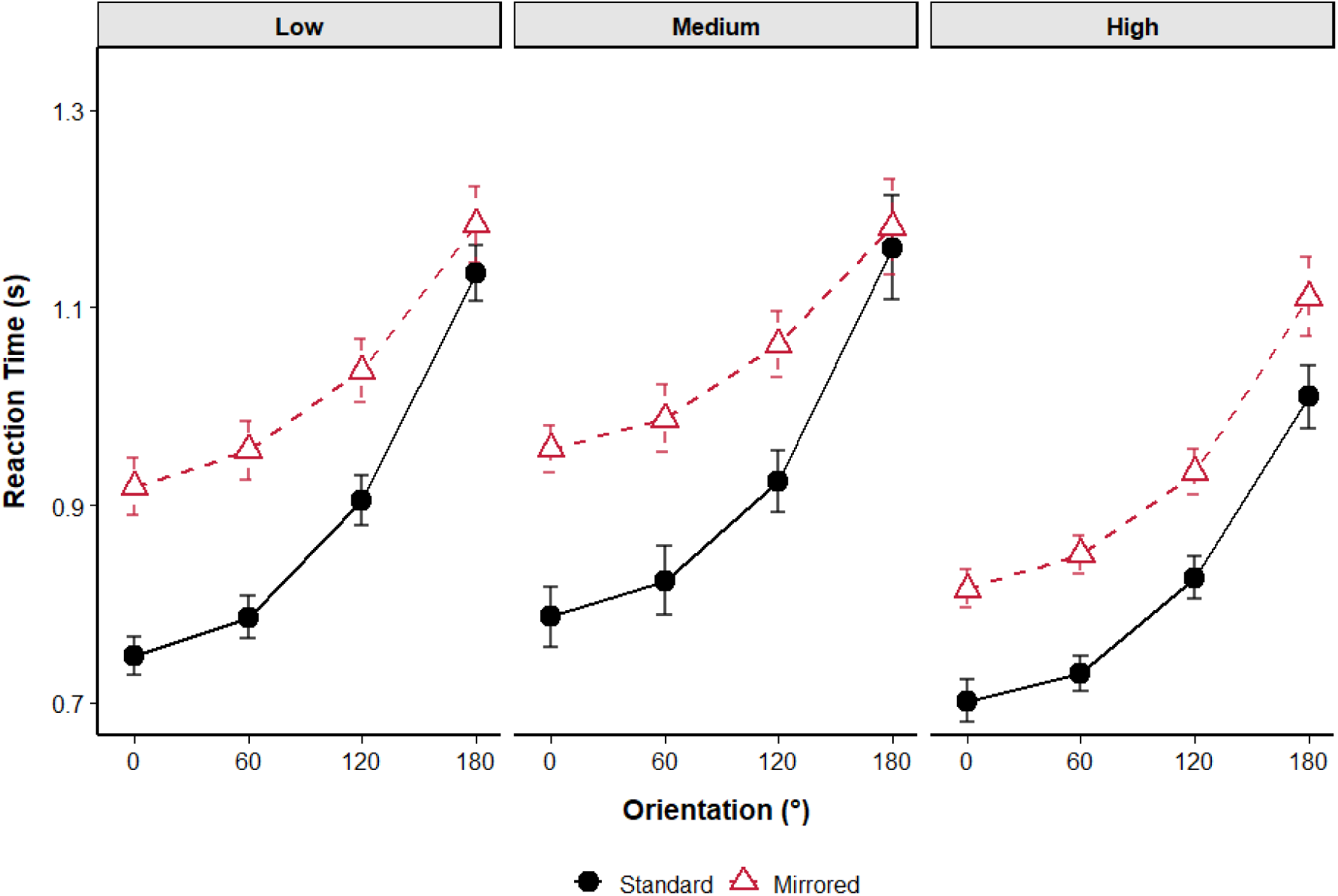
Reaction Time by Version, Orientation and 3D MRT Group. RT increased linearly with rotation angle, and distinctly across versions. Error bars represent ±1 SE.

Accuracy decreased with increasing orientation in both mirrored and standard versions (Figure 6), consistent with greater task difficulty at larger rotation angles. This pattern was further evaluated using a polynomial contrast analysis.

**Figure 6:**
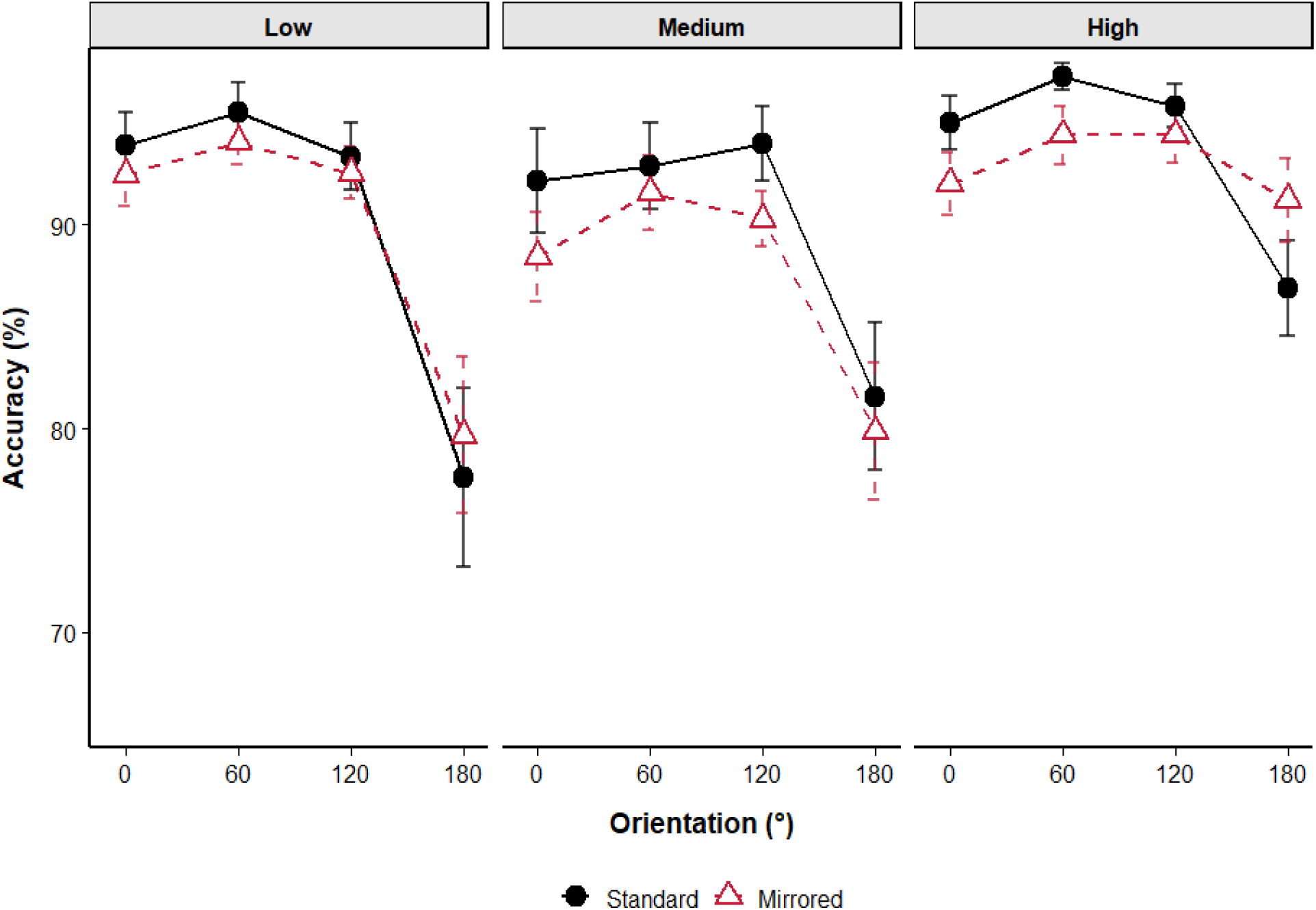
Accuracy by Version, Orientation and MRT Group. High 3D-MR individuals were significantly more accurate than low 3D-MR and Medium 3D-MR individuals especially in the most difficult orientation condition. Low 3D-MR individuals had the steepest drop in accuracy from 120 to 180 degrees rotation.

Orientation effects were significant (linear, t(39) = −5.7; quadratic, t(39) = −7.2; both ps < .001) reflecting that accuracy declined overall as angle increased, but with a curvilinear modulation across orientation levels. The main effect of Parity was not significant at p <.005, nor was its interaction with Orientation effects (highest t(39)= 2.6).

The forward-selected regression model contained just one significant predictor for 3D-MRT score: The linear decline with increasing angle in accuracy (B=.169, r=.41, t(38)=2.7, p<.01). Although adjusted R² square was higher for the 2-predictor model and AIC was lower, BIC was in fact higher, suggesting that the 1-predictor model was more optimal. As can be seen in Figure 6, the forward selected model confirms accuracy declined more gradually across orientations for individuals with higher MRT scores.

### 3.4. ERP: RRN/ P3b

Figure 7 shows the RRN scalp distribution, visualized as the difference between 180° and 0°, with maximal negativity at Pz. Waveforms at CPz are shown in Figure 8A. Note that a specific electrode (CPz) was picked only for visualization; this also holds for the classification into low, medium, and high 3D-MRT scores. Of all the possible combinations of Orientation (0-order, linear, quadratic) and Electrode (0-order, linear, quadratic), only Electrode (linear) and Orientation (linear) x Electrode (linear) were not significant at p < .003. For the other 7 contrasts, all p values were below 0.001, the smallest t(39) value being 3.8. Figure 8B summarizes the pattern of average RRN/ P3b amplitudes per angle (orientation) and per group. Note that, as discussed in the Introduction, the amplitude values in figure 8B reflect both P3b and RRN contributions, with the RRN manifesting as the orientation-dependent decrease in amplitude.

**Figure 7:**
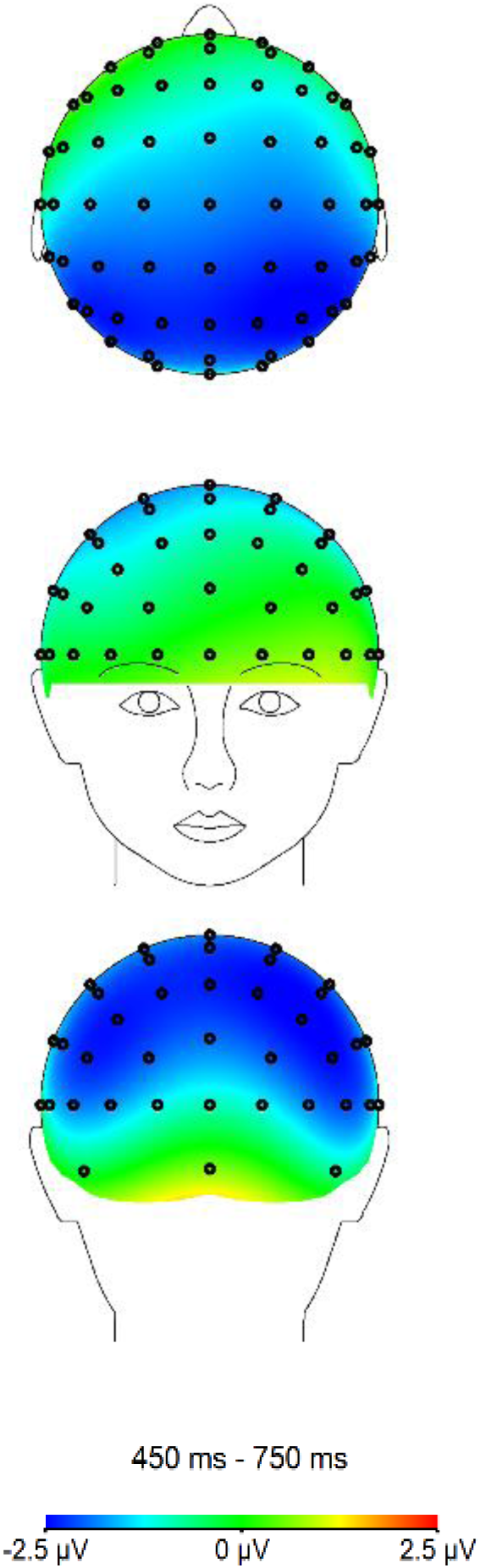
RRN Topography. Voltage difference map (180° - 0°) during the 450-750ms time window, averaged across mirrored and standard parity. The RRN effect (blue = more negative) showed maximal negative amplitude over Pz.

**Figure 8:**
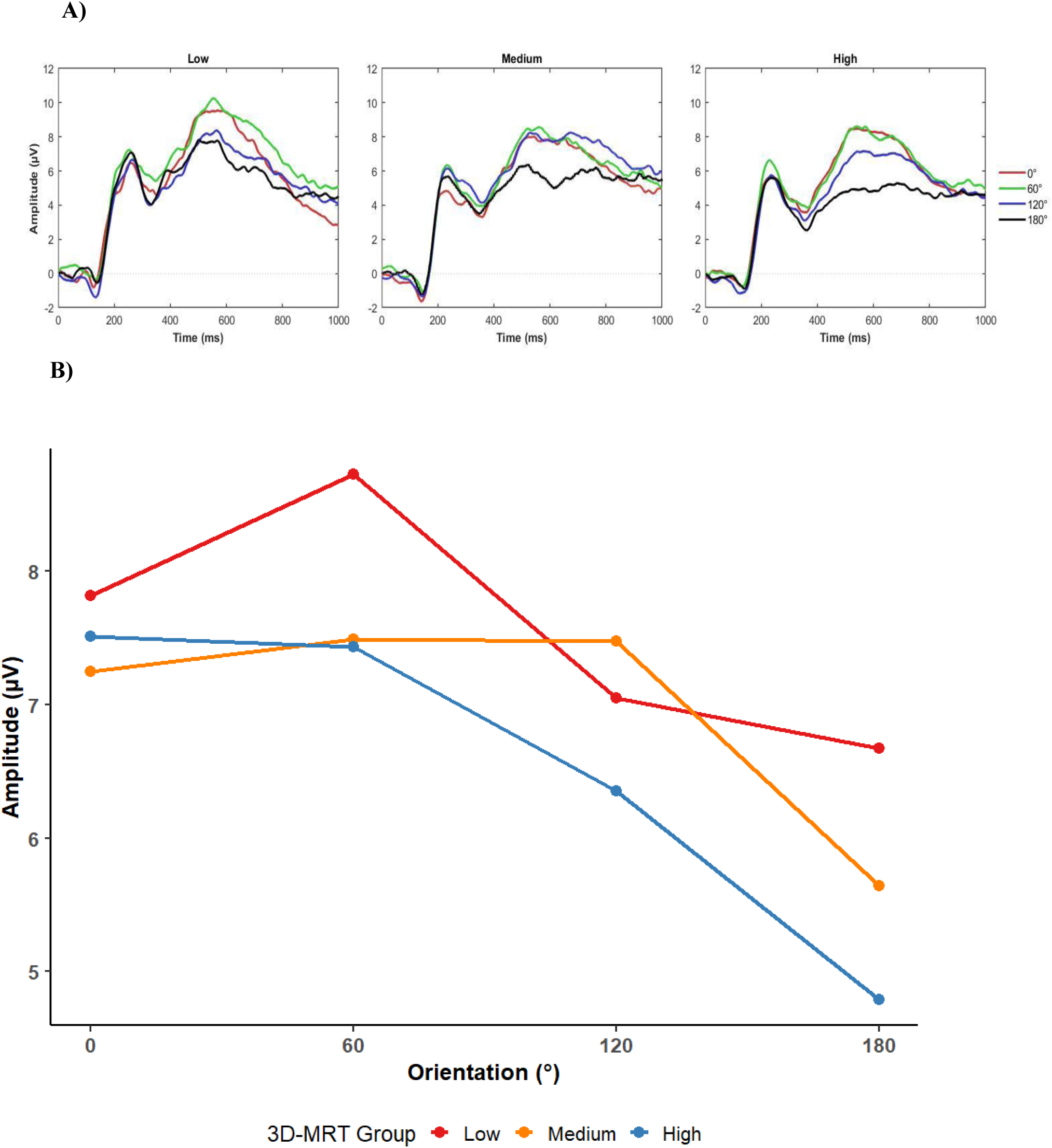
RRN waveforms and slope at CPz by 3D-MRT group. **A)** High 3D-MRT individuals have a more pronounced negativity at 180° degrees in the RRN time window (450-750 ms). **B)** There is a steeper drop (increase in negativity) for the high MRT group relative to the other groups in the most difficult condition at 180°.

Across all participants, amplitudes became progressively more negative as angular disparity increased from 0° to 180°. Both linear and quadratic Orientation trends were significant across all electrodes. The linear RRN trend was strongest for the middle electrodes (CPz and Pz; Orientation (linear) x Electrode (quadratic)), as can be seen in Figure 9. This Orientation effect was slightly biased towards the more anterior electrodes (Orientation (linear) x Electrode (linear)), as was the quadratic trend for Orientation, reflecting that at anterior electrodes, the 180° condition stands out more profoundly from the other conditions. There was no effect of or interaction with the effect of Parity.

**Figure 9.**
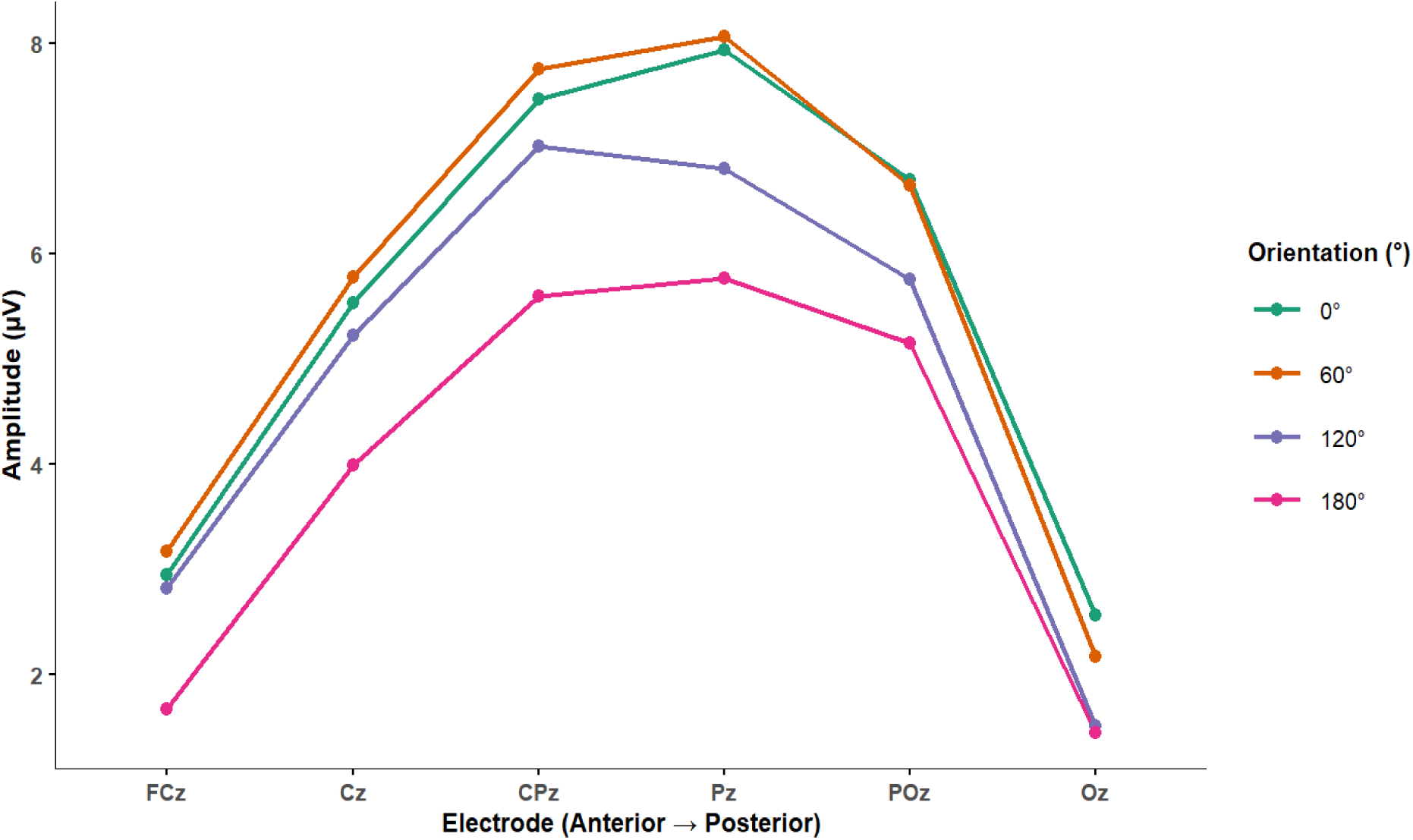
RRN quadratic effect. Increases in angle (linear) results in a more negative mean amplitude across all midline electrodes. This effect is quadratic in that it is more pronounced at the midline electrodes (CPz, Pz).

The forward regression analysis (summarized in Table 1) revealed two significant predictors for 3D-MRT. This was convergingly confirmed by adjusted R^2^, AIC, and BIC values. First, Orientation (linear) x Electrode (quadratic) reflects that high 3D-MRT scores were associated with a stronger linear Orientation trend in particular at the middle electrodes (CPz, Pz). The overall pattern of the linear orientation effect being maximal at central-parietal electrodes is shown in Figure 9. The interaction with 3D-MR ability is illustrated in the waveforms in Figure 8A: individuals with higher 3D-MR ability show more negative amplitudes for larger rotation angles. Figure 8B depicts amplitudes at CPz across rotation angles for low, medium, and high 3D-MRT groups, revealing that high 3D-MR participants exhibited a steeper increase in negative amplitude at higher angles and greater negative amplitude at 180°. The second predictor reflects that, averaged across all rotation angles, a significant quadratic electrode distribution was observed, suggesting individual differences in the spatial distribution of activity during the RRN time window that were independent of orientation. Specifically, low 3D-MRT scorers showed a higher mean amplitude (more positivity) at central-parietal sites (CPz, Pz) relative to medium and high scorers, with group differences being less pronounced at frontal and occipital electrodes, reflecting the quadratic electrode pattern (Figure 10).

**Figure 10.**
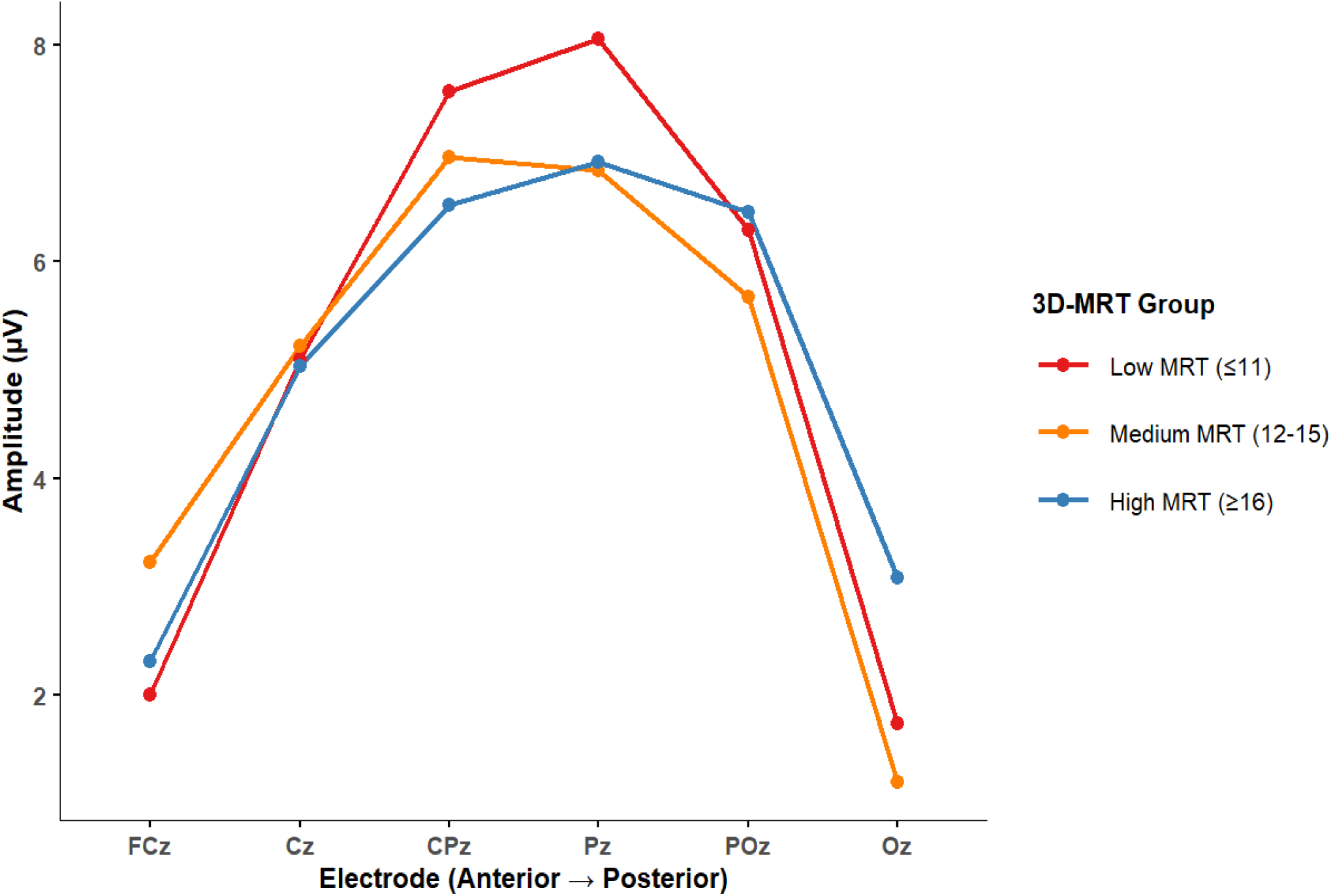
Electrode (quadratic) contrast by 3D-MRT group. Mean amplitude across midline electrodes by 3D-MRT group (orientations and parity collapsed). Low 3D-MRT scorers show higher mean amplitude at CPz and Pz relative to medium and high scorers, suggesting greater reliance on cognitive resources relative to visuospatial representations in lower-ability individuals.

**Table 1.**
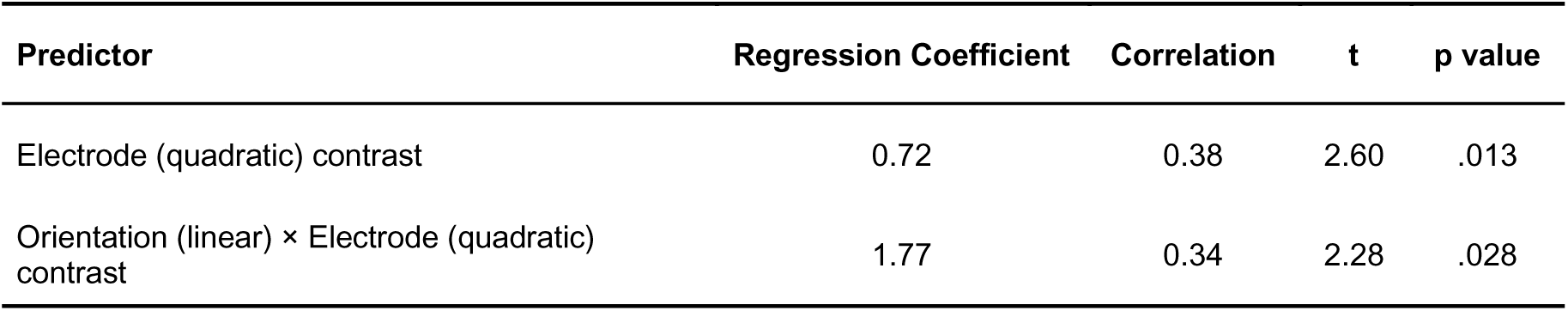
Regression Table. These were the final two significant predictors for 3D-MRT scores resulting from the stepwise regression procedure.

| Predictor | Regression Coefficient | Correlation | t | p value |
| --- | --- | --- | --- | --- |
| Electrode (quadratic) contrast | 0.72 | 0.38 | 2.60 | .013 |
| Orientation (linear) × Electrode (quadratic) contrast | 1.77 | 0.34 | 2.28 | .028 |

### 3.5. ERP: Exploratory Analysis

The exploratory analysis revealed a number of effects that mostly reflect the RRN effects already noted in the dedicated analysis; they are summarized in the Supplementary Material. However, there was one effect outside the RRN window that survived the baseline-cluster criterion. This concerned a main effect of Parity (F(1,39)=21.2; p < .0001).

This effect occurred at a relatively short 200-300-ms latency. Figure 11 characterizes the main features of this effect. The waveforms reveal a discrete deflection that is more positive for mirrored than for standard letters with a maximal effect at F2. In addition, this effect did not significantly correlate with 3D-MRT score (r = −0.11, p = 0.52).

**Figure 11:**
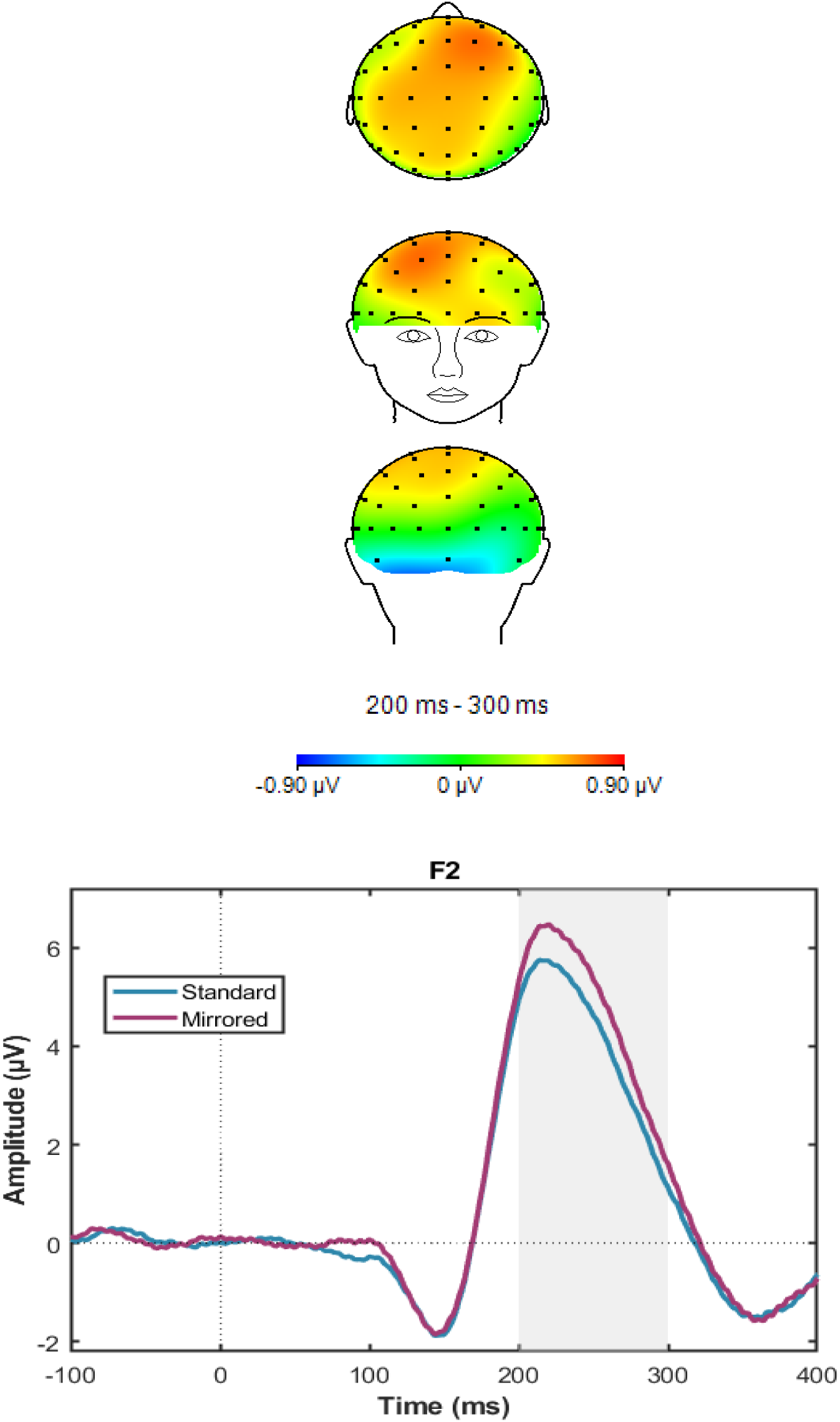
Early Parity effect. Voltage difference map averaged across orientations and MRT (Mirrored - Standard) during the early 200-300 ms time window. The difference is maximal at F2 electrode. Mirrored stimuli are more positive relative to standard stimuli collapsed across all orientations at the time window 200-300 ms.

## 4. Discussion

This study examined how differences in 3D mental rotation capabilities are associated with behavioral and neural markers of 2D mental rotation. Specifically, it was investigated whether individuals with high 3D mental-rotation test (3D-MRT) scores exhibited attenuated behavioral costs as angular disparity increased. Our results show that reaction times increased with angular disparity across both standard and mirrored stimuli, regardless of 3D-MRT score. The mirrored stimuli elicited longer reaction times compared to the standard stimuli especially for small angular disparity. Accuracy decreased with an increase of angular disparity regardless of parity of the stimuli presented (standard/mirrored). Importantly, 3D-MR ability was associated with a steeper drop in accuracy when angular disparity increased for low 3D-MRT scorers. This pattern indicates that higher 3D-MR ability is associated with smaller behavioral costs as rotational demands increase.

Regarding neural measures, the primary prediction was that the rotation-related negativity (RRN) would manifest as an ERP modulation by rotation angle, with increasingly negative amplitudes at larger angular disparities (Farah & Peronnet, 1989; Wijers et al., 1989; Heil et al., 2002; Núñez-Peña & Aznar-Casanova, 2009). This pattern was confirmed in the present data (Figure 8A) and indicates that the current paradigm robustly elicited the characteristic RRN. Regardless of parity, especially in (central-) parietal regions, stimuli rotated to 180° produced the most negative amplitudes in the 450–750 ms window, with 120° yielding an intermediate negativity between 0°-60° and 180° (Figure 9).

In order to determine whether individual differences in spatial ability are associated with the RRN, we examined the relationship between 3D-MRT scores and linear and quadratic RRN slope, in a model that also included orientation-independent ERP values. Higher 3D-MRT scores were associated with a steeper increase in RRN negativity across rotation angles, most pronounced at 180°, relative to lower 3D-MRT groups (see Figure 8). This pattern is consistent with the representation perspective: individuals with stronger 3D-MR ability show greater recruitment of visuospatial representations during rotational transformation, as indexed by the RRN (Heil et al., 2002). This finding is inconsistent with the alternative hypothesis holding that resource allocation dominates over representation recruitment in high scorers, which would have predicted a less negative RRN and stronger P3b for high scorers. Instead, the results support the interpretation that representation recruitment dominates over resource recruitment in high 3D-MRT individuals.

In addition to the rotation-angle effects, 3D-MR ability was associated with differences in the spatial distribution of activity across electrodes, irrespective of angular disparity. Specifically, low 3D-MRT scorers showed a more pronounced central-parietal amplitude distribution compared to high scorers (Figure 10), likely reflecting a greater P3b contribution — indexing greater allocation of cognitive resources — in lower-ability individuals. Together, both findings converge on the same interpretation: high 3D-MRT scorers rely more heavily on visuospatial representations during rotation, while low 3D-MRT scorers show a relatively greater contribution of cognitive resource allocation. This assertion remains partially speculative and could be supported in future work by, for example, estimates of subjectively experienced difficulty, or additional EEG measures such as theta oscillations.

ERP-wise only one new effect of parity (whether the rotated letters were in standard or mirrored orientation) was found in an exploratory analysis. This effect between 200 and 300 ms latency preceded the RRN window and could be suggestive of an non-planar discrimination that begins earlier during early perceptual encoding, consistent with suggestions that mirrored objects require an additional axis flip by which mirrored letters are returned to their canonical orientations (Quan et al., 2020; Zhao et al., 2023). However, this should be interpreted cautiously because it needs replication. For the time being, one further clue is that reaction times were marked by a sub-additive interaction between Parity and Orientation effects. A general explanation for such sub-additive effects invokes parallel processing, in this case of Parity and Orientation (Sternberg, 1969; Pashler, 1984).

Specifically, this “locus-of-slack” logic (e.g., Franz et al., 2008) dictates that the longer reaction times resulting from an increase in difficulty from one manipulation (standard versus mirrored) masks reaction-time lengthening due to another manipulation (increasing angular disparity), as long as the two underlying processes proceed independently. Such an interpretation would be consistent with the present finding of two independent ERP effects of parity and orientation, respectively.

### Limitations

The distinction between high and low 3D-MR ability in this study were derived from the distribution within our sample but may not generalize to other populations or testing contexts. Future work may benefit from combining multiple spatial measures to better characterize the underlying components that represent individual differences in spatial ability. Furthermore, because the task involved planar rotations, the study does not directly resolve whether the observed neural efficiency pattern generalizes to full stereoscopic 3-D rotations involving depth transformations, occlusion, or perspective changes.

### Future Research

Future research should continue to examine the strategies that individuals use when performing mental rotation tasks. Prior work suggests that people differ in whether they rely on holistic transformations, piecemeal comparisons, or feature-based heuristics (Bilge & Taylor, 2017, Gardony et al., 2017, Khooshabeh et al., 2013, Nazareth et al., 2019, Zhao et al., 2018). Understanding the different types of strategies used would help clarify what high 3D-MRT scorers are doing differently from lower scorers and whether such differences are reflected in the EEG signal. This includes determining whether a functional mapping exists between strategy use and the differential reliance on visuospatial representation recruitment versus cognitive resource allocation observed in the present study.

The temporal relationship between planar (standard) and non-planar (mirrored) rotation processes warrants further investigation, particularly regarding their sequential ordering (Zhao et al., 2023). The present findings offer two preliminary clues: a sub-additive interaction between Parity and Orientation in reaction times, suggesting these processes overlap in time, and an early ERP effect of Parity at 200–300ms, preceding the RRN and consistent with non-planar discrimination beginning during perceptual encoding, earlier than the rotation stage itself. Future work applying additive factors logic to reaction times could clarify whether the axis-flip required for mirrored stimuli constitutes a discrete processing stage following planar rotation, or whether it operates in parallel with it (Sternberg, 1969).Ultimately, this line of research has direct applications for the previously mentioned domains such as medicine (including veterinary), architecture, aviation, and general education. Establishing that individual differences exist in spatial processing could also offer a means for more personalized approaches to teaching and learning. Particularly, extended reality technologies could bridge the gap for those lower in mental rotational skills and spatial visualization, and these tools could offer support through targeted 3D visualization and stereoscopic input (Khan et al., 2025, Bogomolova et al, 2020).

### Conclusions

The hypothesis that 2D and 3D mental rotation share underlying spatial processing mechanisms was supported in this experiment. Individuals with high 3D-MRT scores showed clear behavioral advantages in the 2D letter rotation task. At the electrocortical level, high 3D-MRT scorers showed stronger RRN modulation with increasing angular disparity, consistent with greater reliance on visuospatial representation recruitment during rotational transformation. Low 3D-MRT scorers, by contrast, showed a more pronounced central-parietal amplitude distribution irrespective of orientation, consistent with greater reliance on cognitive resource allocation. Together, these results support a common spatial processing architecture underlying both 2D and 3D mental rotation, with individual differences in 3D Mental Rotation ability reflected in the balance between visuospatial representation recruitment and cognitive resource allocation.

## Supporting information

supplementary table and images

