## supplementary table and images for "Rotating Letters in the Mind’s Eye: Behavioral and electro-cortical associations with 3D Mental-Rotation Ability"

**Supplementary Material**

Table S1. Effect clusters longer than corresponding baseline clusters; critical p=.001.

| **Effect nr** | **Effect** | **Latency (ms)** |
| --- | --- | --- |
| 1 | Orilin | 300-700 |
| 2 | Orilin × APqua | 400-800; 1000-1100 |
| 3 | Oriqua × APlin | 200-600; 900-1100 |
| 4 | Oriqua × APqua | 800-900 |
| 5 | Oriqua × APqua × LCRqua | 500-700 |
| 6 | Parity | 200-300 |
| *Orilin = linear Orientation effect; Oriqua = quadratic Orientation effect; APlin = linear Anterior-posterior effect; APqua = quadratic Anterior-posterior effect; LCRlin = linear Left-center-right effect; LCRqua = quadratic Left-center-right effect.* | | |

Effects 1 and 2 reflect the RRN being at maximum over the central-parietal region (see main Figure 9). Effect 3 reflects that the disproportionate increase in negativity from 120 to 180 degrees is larger over frontal areas (also visible in Figure 9; See Figure S1). Effect 4 reflects a late quadratic orientation effect with a quadratic anterior-posterior distribution (800-900ms). Effect 5 is illustrated in Figure S2, showing amplitude across left, center, and right electrode columns separately as a potential indicator of lateralization*.* Effect 6 (Parity) is discussed in the main text.

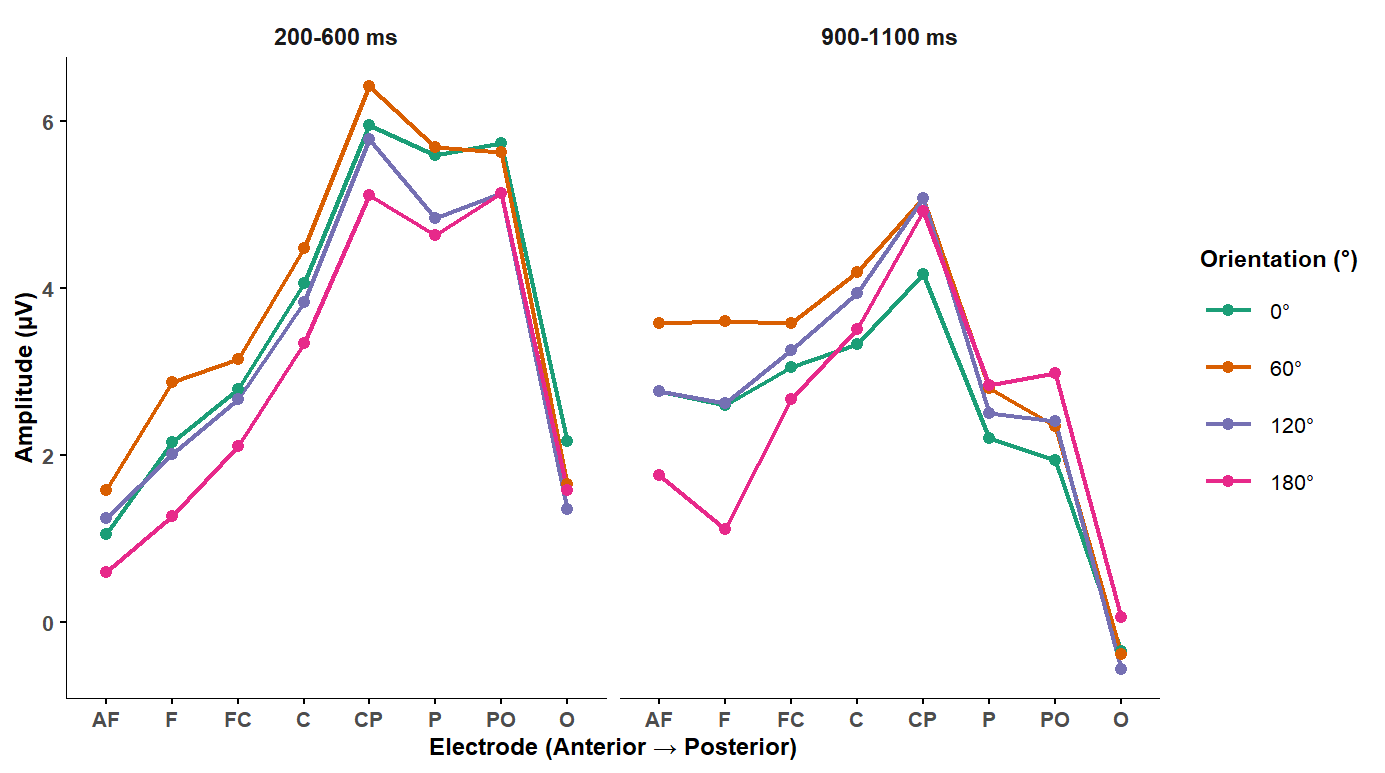

**Figure S1. Mean amplitude across anterior-posterior electrode groups by rotation orientation, for two time windows.** Electrode groups represent the average of left, center, and right electrodes at each anterior-posterior level. The 200-600 ms panel shows an early onset of the linear angle effect with a relatively anterior distribution. The 900-1100 ms panel shows a late re-emergence of orientation-related amplitude differences. Both effects correspond to Effect 3 in Table S1 (Oriqua × APlin).

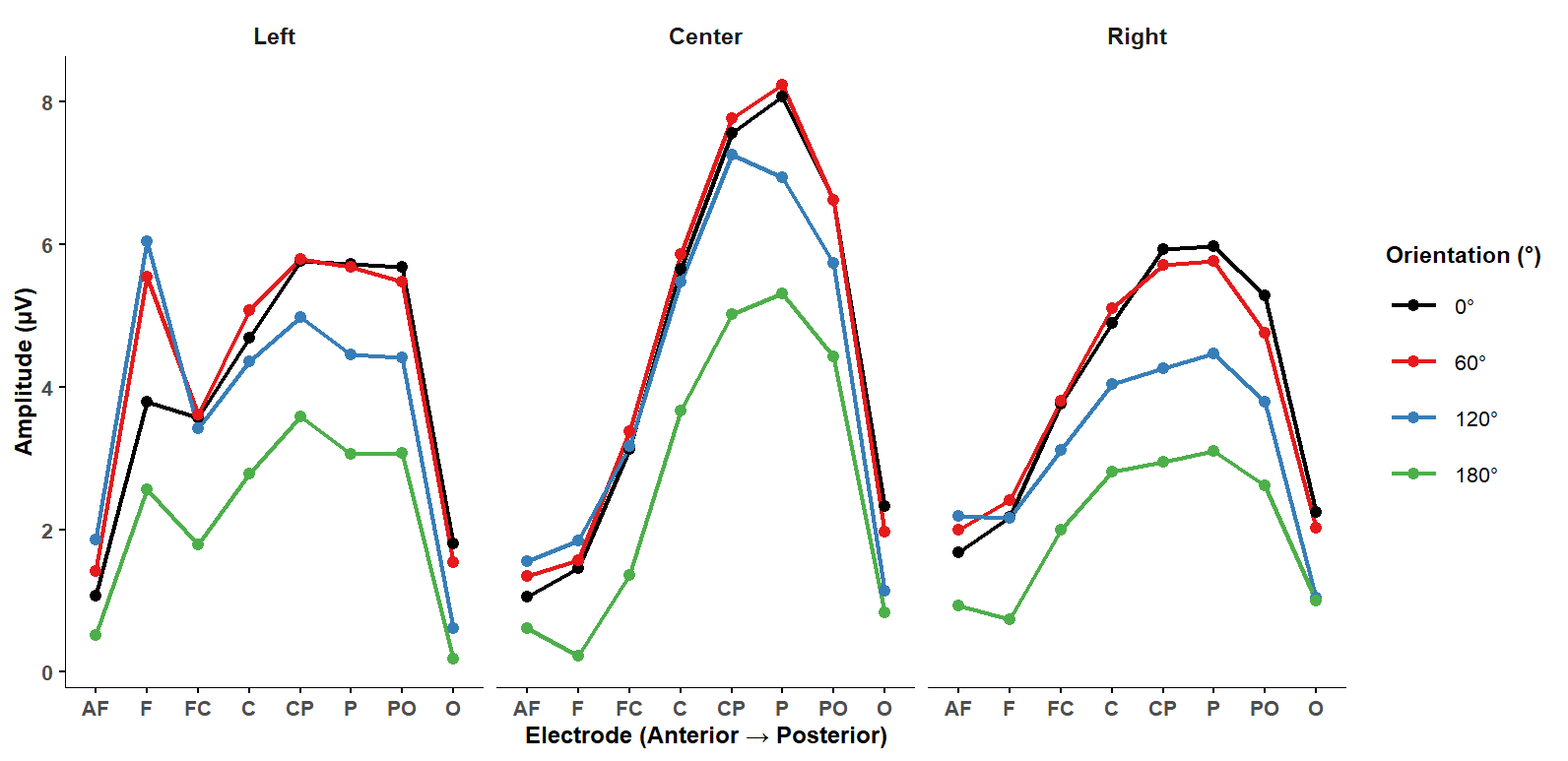

**Figure S2. Mean amplitude across anterior-posterior electrode groups by rotation orientation, shown separately for left, center, and right electrode columns (500-700 ms).** The central-parietal maximum is most pronounced in the center column, consistent with the midline distribution of the RRN. Left and right panels show broadly similar patterns, with subtle differences in the magnitude of the orientation effect across lateral sites, reflecting the Oriqua × APqua × LCRqua interaction (Effect 5 in Table S1).
